# Organic Dust Exposure Induces Stress Response and Mitochondrial Dysfunction in Monocytic Cells

**DOI:** 10.1101/2020.06.30.180323

**Authors:** Sanjana Mahadev-Bhat, Denusha Shrestha, Nyzil Massey, Locke A. Karriker, Anumantha G. Kanthasamy, Chandrashekhar Charavaryamath

## Abstract

Exposure to airborne organic dust (OD), rich in microbial pathogen-associated molecular patterns, has been shown to induce inflammatory responses in the lung resulting in changes in airway structure and function. A common manifestation in lung inflammation is the occurrence of altered mitochondrial structure and bioenergetics, consequently regulating mitochondrial ROS (mROS) and creating a vicious cycle of mitochondrial dysfunction.

The role of mitochondrial dysfunction in airway diseases such as COPD and asthma is well known. However, whether OD exposure induces mitochondrial dysfunction largely remains unknown. Therefore, in this study, we tested a hypothesis that OD exposure induces mitochondrial stress using a human monocytic cell line (THP-1). We examined the mechanisms of organic dust extract (ODE) exposure-induced mitochondrial structural and functional changes in THP-1 cells.

In addition, the effect of co-exposure to ethyl pyruvate (EP), a known anti-inflammatory agent, or mitoapocynin (MA), a mitochondria targeting NOX2 inhibitor was examined. Transmission electron microscopy images showed significant changes in cellular and organelle morphology upon ODE exposure. ODE exposure with and without EP co-treatment increased the mtDNA leakage into the cytosol. Next, ODE exposure increased the PINK1 and Parkin expression, cytoplasmic cytochrome c levels and reduced mitochondrial mass and cell viability, indicating mitophagy. MA treatment was partially protective by decreasing Parkin expression, mtDNA and cytochrome c release and increasing cell viability.

## Introduction

Industrialized agriculture production systems form the backbone of the farm economy in the USA with a large number of workforce and a major contribution to the nation’s GDP (Charavaryamath and Singh, 2006; Nordgren and Charavaryamath, 2018; Sethi et al., 2017). Despite the production efficiency and cheaper price of the food, these industries have occupational hazards in the form of exposure to many on-site contaminants. Among the contaminants, airborne organic dust (OD) and gases (mainly hydrogen sulfide, methane and ammonia), viable bacteria, fungal spores and other microbial products are known to be present (“Respiratory Health Hazards in Agriculture,” 1998). OD comprises of particulate matter (PM) of varying sizes from plant, animal and microbial sources (Vested et al., 2019). Bacterial lipopolysaccharide (LPS) and peptidoglycan (PGN) are the major microbial pattern recognition receptors (PAMPs) present in the OD samples. Agriculture production workers who are exposed to OD report several respiratory symptoms and annual decline in the lung function (Nordgren and Charavaryamath, 2018; Sethi et al., 2017; Wunschel and Poole, 2016). Strategies that reduce the levels of dust in the workplace have shown to have positive health impacts (Senthilselvan et al., 1997).

Respiratory symptoms of exposed workers include bronchitis, coughing, sneezing, chest-tightness, asthma and asthma like symptoms, mucus membrane irritation and other signs. Persistent exposure to OD has been linked to the development of chronic inflammatory conditions, such as chronic obstructive pulmonary disease (COPD) and asthma, including lung tissue damage and decline in lung function (Charavaryamath and Singh, 2006; Wunschel and Poole, 2016). Despite several research groups using both *in vitro* and *in vivo* models of OD exposure, precise cellular and molecular mechanisms leading to chronic lung disease remain largely unknown.

In response to this, an understanding of the mechanisms of induction of airway inflammation is essential as it promotes the development of strategies for the maintenance of lung homeostasis by preserving the balance between pro-inflammatory and anti-inflammatory responses. Studies have shown that OD-mediated lung inflammation is typically characterized by airway hyperresponsiveness (AHR), tissue remodeling, and increased influx of inflammatory cells, particularly neutrophils and macrophages, in lung tissues (Charavaryamath et al., 2005; Sahlander et al., 2012; Sethi et al., 2017). In previous studies we have shown exposure of human bronchial epithelial cells to OD results in the production of reactive oxygen species (ROS), reactive nitrogen species (RNS) and a myriad of pro-inflammatory cytokines such as interleukins IL-1B, IL-6, and IL-8 (Bhat et al., 2019; Nath Neerukonda et al., 2018). Based on evidence, it is also becoming increasingly clear that, in addition to the above factors, abnormal mitochondrial signatures and mitochondrial dysfunction contribute to the pathological mechanisms of lung disease (Cloonan and Choi, 2016). Several *in vitro* and *in vivo* studies have demonstrated the elevation of key enzymes involved in the production of ROS and RNS due to mitochondrial impairment in various inflammatory conditions (Cloonan and Choi, 2012; Eisner et al., 2018; Zhang et al., 2010). Collectively, these findings suggest that targeting multiple pathogenic mechanisms, including mitochondrial impairment, oxidative stress, and other inflammatory processes, could provide an advantage over targeting a single disease pathway.

Mitochondria are dynamic double membraned organelles that possess their own genome and proteome. These are ubiquitously present and are critical for many of the body’s “housekeeping” functions, including synthesis and catabolism of metabolites, calcium regulation, and most importantly generation of ATP by oxidative phosphorylation (OXPHOS) (Tilokani et al., 2018). Whilst the participation of mitochondria in OXPHOS, stress responses and programmed cell death pathways have been well studied over the past decade, the role of mitochondria in the activation and control of the immune response has been of interest. During inflammation, mitochondria can become damaged or dysfunctional leading to impaired cellular respiration and cell death. The presence of dysfunctional mitochondria can lead to oxidative stress which acts as a potent stimulus for exacerbating inflammation (Cloonan and Choi, 2012; Eisner et al., 2018).

The adverse effects of inflammation on mitochondria can be abrogated by several mechanisms. These include the induction of anti-inflammatory responses and antioxidant defenses, maintenance of mitochondrial integrity through the selective removal of dysfunctional mitochondria (mitophagy), and the generation of new organelles to replace them (mitochondrial biogenesis) (Eisner et al., 2018). However, the integration of these compensatory responses, and the interaction between mitochondria and host cells following OD exposure, are not well understood. In order to study these processes, we assessed mitochondrial functions, biogenesis, and mitophagy on exposure to OD alone and in the presence of antioxidant therapies, such as ethyl pyruvate (EP) and mitoapocynin (MA), which have previously been shown to have significant antioxidative functions.

The protective effects of EP have been attributed to its anti-inflammatory, antioxidative and antiapoptotic action. Previously we have shown the effectiveness of ethyl pyruvate (EP) as a non-specific inhibitor of inflammatory cytokine-like high mobility group box 1 (HMGB1) release into the extracellular space in bronchial epithelial cells (Bhat et al., 2019). We also demonstrated that EP downregulates reactive oxygen species (ROS) generation and augmented IL-10 production thus promoting anti-inflammatory effects. Similar results have been shown in LPS injected and ischemic animal models (Venkataraman et al., 2002; Yu et al., 2005). The anti-inflammatory property of EP has been attributed to the inhibition of ROS-dependent signal transducer and activator of transcription (STAT) signaling (Kim et al., 2008).

In addition to using EP, we also tested the efficacy of MA in a OD-induced inflammatory model. In previous studies, apocynin, a plant derived antioxidant, has been used as an efficient inhibitor of NADPH-oxidase complex in many experimental models involving phagocytic and nonphagocytic cells (Stefanska and Pawliczak, 2008). In this study we used triphenylphosphonium (TPP) conjugated apocynin (mitoapocynin, MA) designed to enhance their cellular uptake and target the mitochondria. In contrast to other popular antioxidant therapies, MA has been shown to attenuate ROS and/or RNS generation in both *in vitro* and *in vivo* models of neuroinflammation. In an MPTP-induced neuroinflammatory model, MA treatment was shown to suppress iNOS and various pro-inflammatory cytokines. In addition, MA was shown to inhibit NOX2 activity and reduce oxidative stress (Ghosh et al., 2016; Langley et al., 2017).

In this study we used an immortalized human monocytic cell line (THP1) and tested a hypothesis that OD-exposure induces mitochondrial stress. We further examined whether there is an induction of antioxidant defenses, changes in mitophagy and mitochondrial biogenesis in THP1 cells following exposure to OD in the presence of both a mitochondrial specific NOX2 inhibitor (MA) and an inhibitor of HMGB1 translocation (EP), leading to the maintenance of cellular viability and mitochondrial integrity. Here we demonstrate that mitochondrial specific or general antioxidant therapy, through inhibition of HMGB1 translocation, are vital to cellular recovery following exposure to OD.

## Materials and Methods

### Chemicals and reagents

We purchased RPMI 1640, L- glutamine, penicillin, streptomycin, MitoTracker green, and MitoSOX Red stains from Invitrogen (ThermoFisher Scientific) and fetal bovine serum (FBS) was purchased from Atlanta Biologicals. Antibodies for mitofusins (MFN1/2), DRP1, PINK1, Parkin, OPA1, BNIP3, Cytochrome C, COX4i2, Bcl-2, Bcl-XL, mtTFA, Caspase 1 and Caspase 3 was purchased from Santa Cruz Biotechnology. The anti-HMGB1 antibody, β-Actin antibody and Rhod-2AM dye were obtained from Abcam. MitoApocynin-C_11_ (MA) was procured from Dr. Balaraman Kalyanaraman (Medical College of Wisconsin, Milwaukee, WI), stock solution (10 mM in DMSO) prepared by shaking vigorously and stored at −20°C. MA was used (10 μM) as one of the co-treatments. Ethyl pyruvate (EP) purchased from Santa Cruz Biotechnology, was reconstituted in Ringer’s solution (Sigma-Aldrich) and used at a final concentration of 2.5 μM in the cell culture medium.

### Organic dust extract preparation

Aqueous organic dust extract (ODE) was collected and prepared as previously described (Bhat et al., 2019; Romberger et al., 2002). Briefly, settled surface dust samples from swine housing facilities were collected and 1 g was placed into sterile Hank’s Balanced Salt Solution (10 ml; Gibco). Solution was incubated for one hour at room temperature, centrifuged for 20 min at 1365 x g, and the final supernatant was filter sterilized (0.22 μm), a process that also removes coarse particles. Stock (100%) ODE aliquots frozen at −20°C until use in experiments. The filter sterilized organic dust extract (ODE) samples were considered 100% and diluted to 1-5% (v/v) before use in experiments.

### Cell culture and treatments

Immortalized human monocytic cells (THP1, ATCC TIB-202™) were used in this study. These cell lines have previously been used to study innate inflammatory responses to ODE (Nath Neerukonda et al., 2018). THP1 cells were cultured in RPMI 1640 at 37°C in a humidified chamber with 5% CO_2_. The RPMI 1640 medium was supplemented with 10% (v/v) heat-inactivated FBS, 2 mM L-glutamine, 10 mM HEPES, 1.5 g/L sodium bicarbonate, 1 mM sodium pyruvate, 100 IU/ml penicillin, and 100 μg/ml streptomycin (Gibco) and 1 μg/mL of Amphotericin B (Sigma-Aldrich). Cells were subcultured once a week and the morphology was observed. Approximately 4 to 5-day old cultures were used for experiments. Treatments were done in 1% FBS-containing medium for 24 hours. All groups with treatment details are outlined in figure 1.

**Figure 1.**
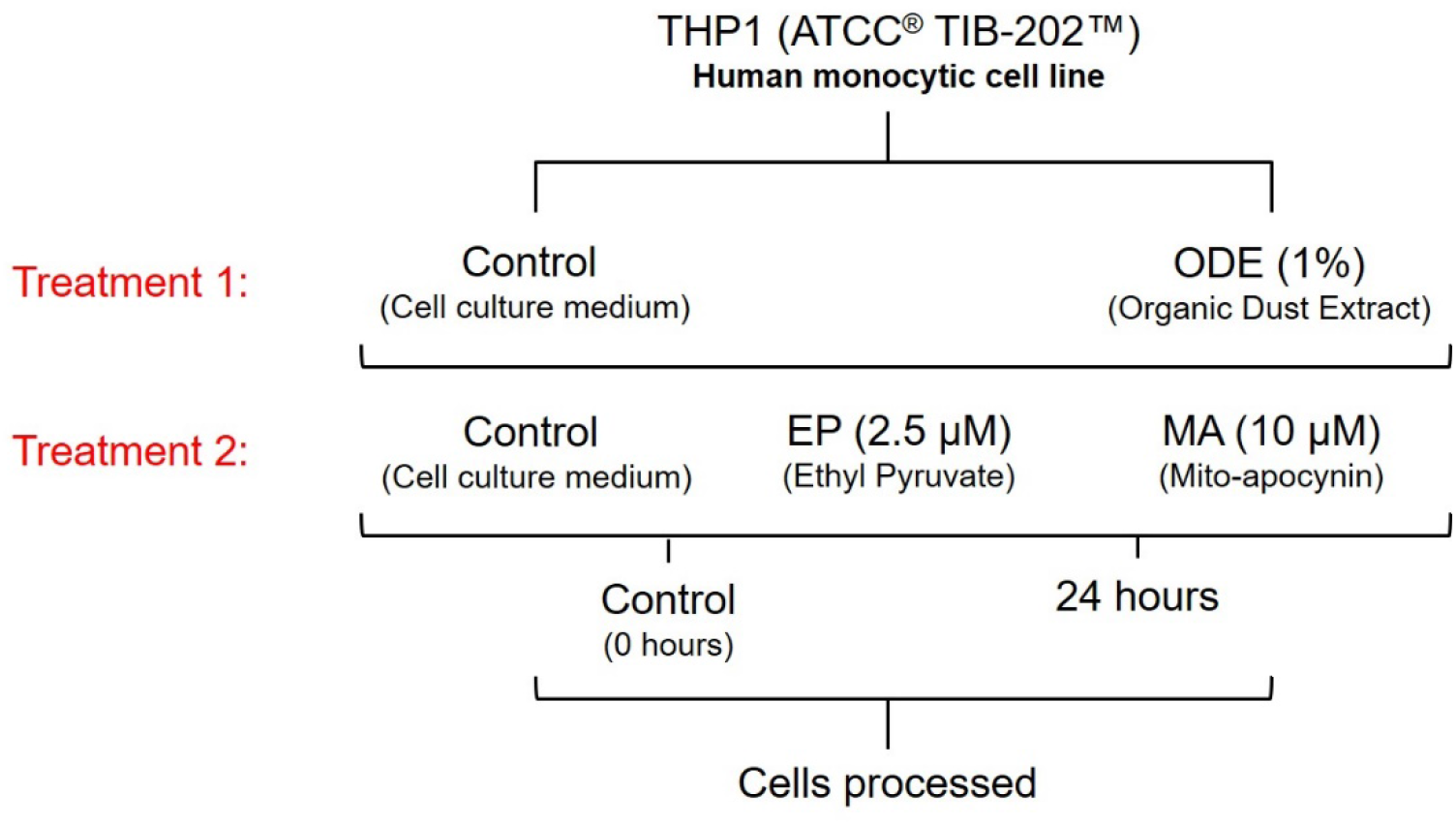
ODE exposure of THP1 cells and antioxidant treatment. THP1 cells were treated with either media (control) or ODE (treatment 1) followed by either media, EP or MA (treatment 2). Cells were processed for various assays at 0 (control), and 24 hours.

Ethyl pyruvate (EP) was reconstituted in Ringer’s solution and used at a final concentration of 2.5 μM in the cell culture medium. Mitochondrial specific drug mitoapocynin (MA) was diluted in dimethyl sulfoxide (DMSO) and used at a final concentration of 10 μM (Ghosh et al., 2016; Langley et al., 2017).

Cells were treated with either medium (control) or ODE (1% v/v) followed by a co-treatment with either EP (2.5 μM) or MA (10 μM) for 24 hours, with corresponding time matched controls. Following the treatments, samples were processed at 24 hours for various assays.

**Table 1.**
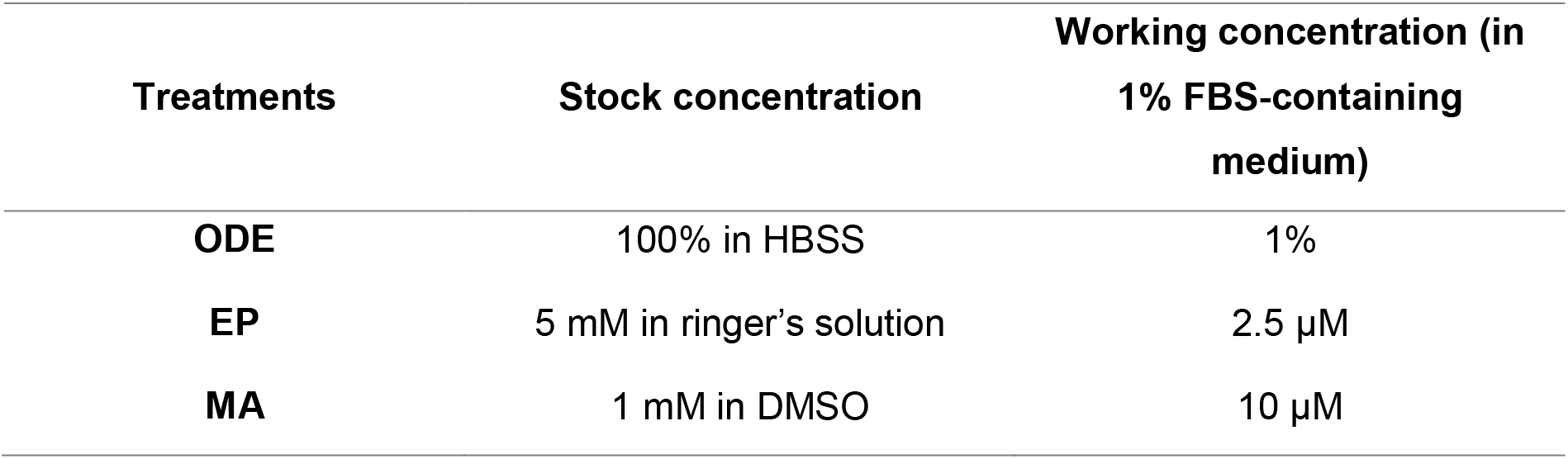
Stock and working concentrations of treatments.

### Cell viability and MTT assay

Prior to conducting experiments, cell viability was assessed. Live/dead cell count was determined by 4% trypan blue dye (EMD Millipore) exclusion and percentage viability was calculated. Population of cells with more than 95% viability were used for the experiments.

The MTT assay has been widely used in the estimation of LC50 and cell viability by measuring the formazan produced when mitochondrial dehydrogenase enzymes cleave the tetrazolium ring (Latchoumycandane et al., 2005). In this study, we used the MTT assay to determine the LC50 of ODE in THP1 cells. Cells were seeded (20,000 cells/well) in a 96-well culture plate and treated with ODE for 24 hours in 1% FBS-containing RPMI medium. After the treatment, the cells were washed with PBS and incubated with 0.5 mg/mL of MTT in 1% FBS-containing RPMI medium for 3 hours at 37°C. The supernatant was removed, and MTT crystals were solubilized in 100 μl of DMSO. Absorbance was measured with the SpectraMax spectrophotometer (Molecular Devices Corporation) at 570 nm with the reference wavelength at 630 nm.

### Transmission Electron Microscopy

Post-treatment THP1 cells were washed twice with RPMI and fixed for 45 min in a fixative solution (2% Glutaraldehyde in complete culture medium). The samples were centrifuged, and the pellet fixed again with 1.5% Glutaraldehyde solution in Na-Cacodylate buffer 0.1 M. A final post-fixation (2 h) in 1% OsO_4_ solution in Na-Cacodylate buffer 0.1 M was performed. The samples were mixed with uranyl acetate 2% (w/v) and incubated for 5 min, and then, 5 μl was applied to carbon-coated copper grids. Images were taken using a JEOL 2100 200-kV scanning and transmission electron microscope with a Thermo Fisher Noran System 6 elemental analysis system. TEM was operated at 80 kV, and images were obtained at 2,000× to 12,000× magnification. (Electron Microscopy Facility, Iowa State University).

### Morphological analysis

Mitochondrial shape descriptors and size measurements were obtained using ImageJ (National Institutes of Health) by manually tracing only clearly discernible outlines of mitochondria on TEM micro-graphs (Picard et al., 2013). Surface area (or mitochondrial size) is reported in μm^2^; perimeter in μm; circularity [4*(surface area/perimeter2)]; and Feret’s diameter represents the longest distance (μm) between any two points within a given mitochondrion. Computed values were imported into Microsoft Excel and Prism 8.0 for data analysis.

### Subcellular fractionation

Whole cell and subcellular protein lysate extractions (cytosol and mitochondria) were performed at 4°C using cold reagents. For whole cell protein lysates, cell pellets were subjected to lysis using RIPA buffer [with protease and phosphatase inhibitors] (ThermoFisher Scientific). Subcellular fractionation of cell pellets for isolation of mitochondria was done using the Mitochondria Isolation Kit for Cultured Cells (ThermoFisher Scientific) according to the manufacturer’s instructions. The whole cells, cytosolic fraction and isolated mitochondria were lysed with RIPA buffer [with protease and phosphatase inhibitors] for 30 min at 4°C and periodic sonication on ice, followed by centrifugation to collect lysate. Protein concentration of fractions were determined by Bradford assay (Bio-Rad) and were stored at −80°C until use.

### Mt DNA isolation and long-range PCR

To determine mitochondrial DNA (mtDNA) leakage, mtDNA was isolated from mitochondria-free cytosolic fraction of the cells. Mitochondrial DNA from cytosolic fractions was extracted using the Genomic DNA Purification kit (ThermoFisher Scientific) as per the manufacturer’s instructions. The purity and concentration of the isolated DNA was measured using NanoDrop (NanoVue Plus Spectrophotometer, GE Healthcare). Due to low concentrations, the mtDNA was first amplified by long range PCR. The primers used were: mtDNA gene, sense: 5′-TGAGGCCAAATATCATTCTGAGGGGC-3’ and antisense: 5′-TTTCATCATGCGGAGATGTTGGATGG-3’ (Liu et al., 2015). PCR reactions were performed at 94°C for 1 min followed by 30 cycles at 98°C for 10 s, 60°C for 40 s, 68°C for 16 min and a final elongation for 10 min (Liu et al., 2015). Confirmation of the presence of mtDNA was done by separating the product by electrophoresis on a 0.8% agarose gel stained with ethidium bromide. The concentration of amplified mtDNA obtained was adjusted to ensure equal amounts of template mtDNA in each sample used for qPCR reaction.

### Quantitative Real-Time PCR

Change in fold change of mtDNA was measured by qPCR with primers specific to mitochondrial NADH dehydrogenase 1 *(mtND1).* 5 μL of SYBR Green Mastermix (Thermo Fisher Scientific), 1 μL of primers, 2-3 μL of DNase/RNase free water and 1 μg of amplified mtDNA was used. The primers for genes of interest were synthesized at Iowa State University’s DNA Facility. The primers used were: mtND1 gene, sense: 5’-GGCTATATACAACTACGCAAAGGC-3’ and antisense: 5’-GGTAGATGTGGCGGGTTTTAGG-3’; 16s (housekeeping gene), sense: 5’-CCGCAAGGGAAAGATGAAAGAC-3’ and anti-sense: 5’-TCGTTTGGTTTCGGGGTTTC-3’. No-template and no-primer controls and dissociation curves were run for all reactions to exclude cross-contamination. The qRT-PCR reactions were run in a QuantiStudio 3 system (ThermoFisher) and the data was analyzed using 2^−ΔΔCT^ method (Livak and Schmittgen, 2001).

### Western blot analysis

Lysates (whole cell, cytosol and MT) containing equal amounts of protein (20 μg/sample), along with a molecular weight marker (Bio-Rad), were run on 10–15% sodium dodecyl sulfate/polyacrylamide gel electrophoresis (SDS-PAGE) as previously described(Bhat et al., 2019). Proteins were transferred to a nitrocellulose membrane and nonspecific binding sites were blocked with Licor Odyssey blocking buffer. To investigate mitochondrial dysfunction, the membranes were then incubated with different primary antibodies such as MFN1, MFN2, OPA1, DRP1, PINK1, Parkin, BNIP3, Cytochrome C, COX4i2, Bcl-2, Bcl-XL, mtTFA, SOD2, Caspase 1 and Caspase 3 (1:1000 dilution). HMGB1 expression in mitochondrial fractions was measured using anti-HMGB1 antibody (1:5000 dilution). Next, membranes were incubated with one of the following secondary antibodies: Alexa Fluor 680 goat anti-mouse, Alexa Fluor 680 donkey anti-rabbit or Alexa Fluor 800 donkey anti-rabbit (1:10,000; Invitrogen). To confirm equal protein loading, blots were probed with a β-actin antibody (AbCam; 1:10,000 dilution). Western blot images were captured using Odyssey® CLx IR imaging system (LI-COR Biotechnology) and analysis was performed using ImageJ (National Institutes of Health).

### Mitochondrial activity and MitoSOX assay

Cells were seeded (50,000 cells/well) in a 96-well culture plate and treated for 24 hours. After treatment, the media was removed and 100 μl of 200 nM MitoTracker green and 5 μM MitoSOX red dye diluted in 1% FBS-containing RPMI medium was added into each well and incubated at 37°C for 15 min. Following incubation, the cells were washed with 1% FBS-containing RPMI medium and fluorescence intensity was measured by spectrophotometer reading taken at excitation/emission wavelengths of 485/520 nm and 510/580, respectively (SpectraMax M2 Gemini Molecular Device Microplate Reader). The results were expressed as percentage mean fluorescence intensity (%MFI) relative to control.

### Mitochondrial calcium influx measurement by rhod-2AM staining

Mitochondrial calcium influx ([Ca^2+^]_mito_) in THP1 cells was measured using the rhod-2AM dye. The concentration of the isolated mitochondria was measured by Bradford assay in order to maintain consistency in the number of mitochondria loaded into the wells of a 96-well plate. A protein concentration of 100 ug was loaded into each well and10μM Rhod-2AM (Abcam) dye diluted in 1% FBS-containing RPMI medium was added and incubated at 37°C for 30 minutes in order to stain the mitochondria. The cells were washed with 1% FBS-containing RPMI medium and fluorescence was read at excitation/emission wavelengths of 552 nm/581 nm using a spectrophotometer reader (SpectraMax M2 Gemini Molecular Device Microplate Reader).

### Griess assay

Griess assay was performed as described previously (Gordon et al., 2011). Briefly, nitric oxide secretion was measured (representing reactive nitrogen species (RNS)) as nitrite levels in cell culture media using Griess reagent (Sigma-Aldrich) and sodium nitrite standard curve, prepared using a stock solution of 200 μM. The assay was performed in a 96 well-plate and absorbance was measured at 550 nm (SpectraMax M2 Gemini Molecular Device Microplate Reader). The results were expressed as μM concentration of nitrite secreted.

### Statistical analysis

Data analysis and graphical representation was performed using GraphPad Prism 8.0 software (GraphPad Prism 8.0, La Jolla, CA, USA). Data was analyzed with one-way ANOVA with Tukey’s multiple comparison test and a p-value of < 0.05 was considered to be statistically significant.

## Results

### Exposure to ODE impacts the cellular and mitochondrial morphology

TEM images showed that THP1 cells treated with media alone (controls) showed normal morphology with healthy mitochondria (Fig. 2a & 2b). After ODE treatment, cytoplasmic vacuolization and pseudopod formation was observed which suggests differentiation of the cells to form activated macrophages (Fig. 2a) (Krysko et al., 2006). In addition, the mitochondria seemed larger in size and some were elongated with reduced cristae number and/or deformed cristae (Fig. 2b). On addition of 2.5 μM ethyl pyruvate (EP), formation of multinucleated giant cells was observed, which is stated to be commonly observed in diverse infectious and non-infectious inflammatory conditions (Milde et al., 2015; Miron and Bosshardt, 2017). Similar to ODE exposure, the mitochondria were swollen and showed disorganized cristae, along with the presence of calcium sequestration bodies in the mitochondrial matrix (Fig. 2b). In contrast, exposure to 10 μM mitapocyanin (MA) seemed to oppose the impact of ODE on the cells and restore it (Fig. 2b). Almost no cytoplasmic vacuolization was observed, and mitochondria showed decreased signs of damaged cristae, albeit conformed to an elongated morphology (Fig. 2a & 2b). This suggests that MA could have a protective effect on ODE exposed macrophages.

**Figure 2.**
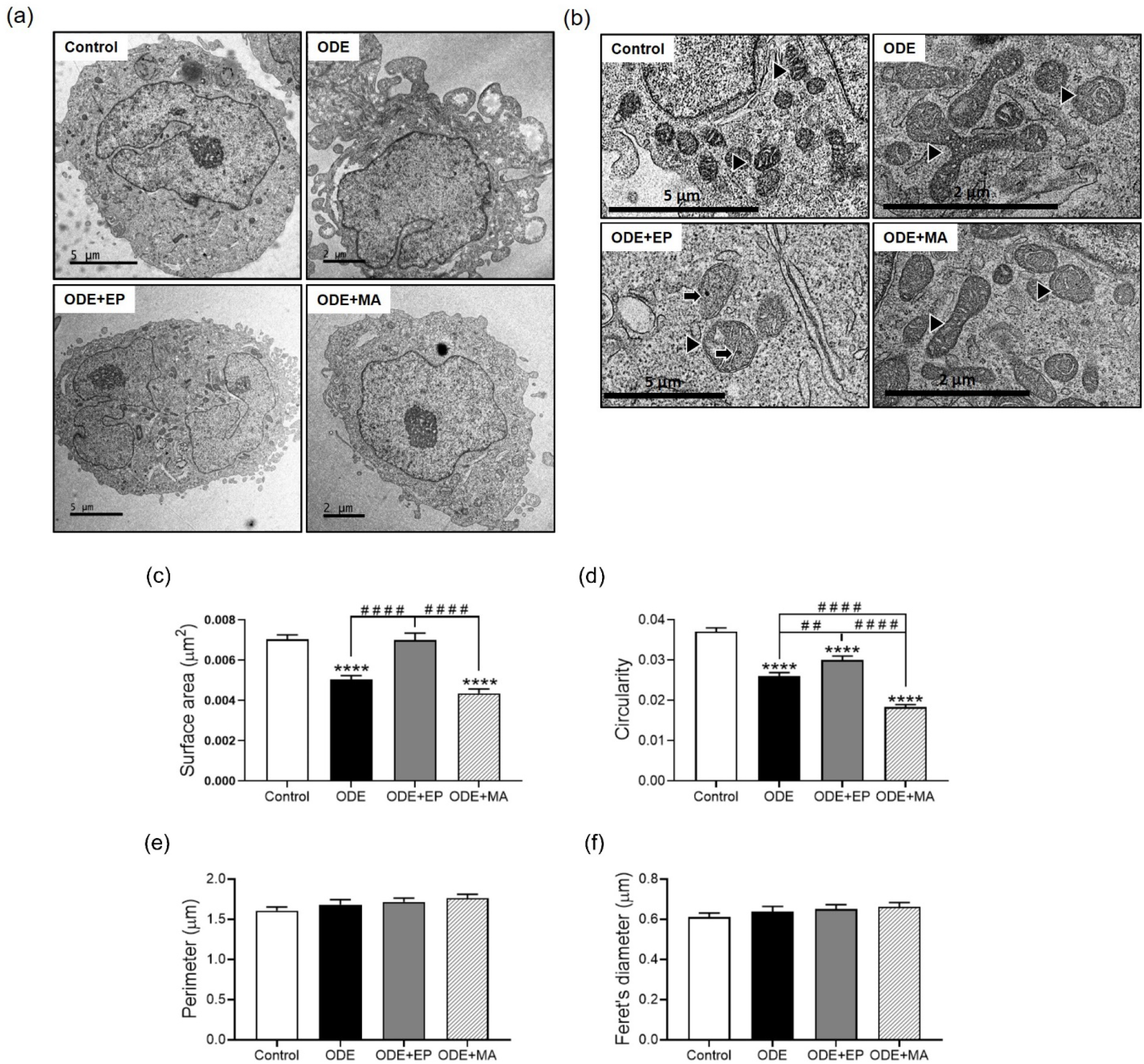
ODE exposure induces differentiation of THP1 cells and changes in mitochondrial morphology. Transmission electron microscopy (TEM) of THP1 cells treated with ODE and antioxidant therapy for 24 hours shows changes in cellular and mitochondrial morphology at the ultrastructural level. Compared to controls, cells undergo differentiation into activated macrophages, with increased vacuolation and pseudopod formation on treatment with ODE (1%; a). Scale bar, 2-5 μm. A number of mitochondria show changes in morphology (fission/fusion) and swelling (b), along with presence of calcium sequestration bodies within the mitochondrial matrix in cells co-treated with 2.5 μM of EP, and noticeably healthier mitochondria with some morphological changes (fission/fusion) in cells co-treated with 10 μM of MA. Morphological parameters of mitochondria on treatment was analyzed by ImageJ (c-f). Significant change in the surface area (c) and circularity (d) indicative of mitochondrial fragmentation, and no change in perimeter (e) and the feret’s diameter (f). Data analyzed via one-way ANOVA with Tukey’s multiple comparison test, ^# or^ *p < 0.05, ^# # or^ **p < 0.01, ^# # # or^ ***p < 0.001, ^# # # # or^ ****p < 0.001 and are represented as Mean ± SEM with n = 126 mitochondria/treatment.

In order to quantify what was observed in the TEM images, mitochondria were individually traced from the TEM. Compared to controls, exposure to ODE significantly reduced the mitochondrial size (Fig. 2c). A similar decrease was seen in the presence of MA as well, whereas EP significantly increased the size, similar to that of control (Fig. 2c). A similar pattern was seen with the mitochondrial circularity, co-treatment with MA showing the most decrease compared to other treatments (Fig. 2d). Other morphological parameters such as perimeter, and Feret’s diameter did not differ between any of the treatment conditions (Fig. 2e & 2f). Taken together, this suggests that exposure to ODE activates a specific mechanism that acts towards altering the mitochondrial dynamics within the cell.

### Targeted antioxidant therapy promotes mitochondrial fission

The mitochondrial membrane is remodeled continuously through cycles of fission and fusion events. The delicate balance of these events helps in controlling mitochondrial structure and function (Tilokani et al., 2018; Wai and Langer, 2016). In order to accurately interpret the impact of ODE exposure on the morphology of mitochondria, the expression of markers responsible for these dynamic events was observed. During ODE exposure, the expression of mitofusin 2 (MFN2) was significantly increased compared to control (Fig. 3a & 3c). In contrast, mitofusin 1 (MFN1) and optic atrophy 1 (OPA1) did not show any significant changes in protein levels between control and treatments (Fig. 3a, 3b & 3d). MFN1 and MFN2 are outer mitochondrial membrane GTPases that are responsible for the promotion of mitochondrial fusion (Tilokani et al., 2018). MFN2 alone can induce mitochondrial fragmentation and is a crucial regulator of mitochondria-endoplasmic reticulum (ER) contact site tethering (Filadi et al., 2018; Tilokani et al., 2018). Increased expression of dynamin-related protein 1 (DRP1) was observed in both ODE and MA treatments, indicative of mitochondrial fission as a result of oxidative stress and/or increased cytosolic calcium levels (Fig. 3e) (Eisner et al., 2018). Furthermore, to corroborate whether mitochondrial morphological changes are associated with mitochondrial number, change in mitochondrial mass on treatment was measured by mitotracker green fluorescence. There was a significant increase in the mitochondrial mass on exposure to ODE and EP, while on exposure to MA levels were comparable to controls (Fig. 3f). This could be a cellular response in order to compensate for the reduced mitochondrial function (Nugent et al., 2007). Collectively, the results suggest an effort to rescue mitochondrial biogenesis by increased MFN2 mediated fusion in response to ODE-induced cellular stress by maintaining a functional population of mitochondria within the cell.

**Figure 3.**
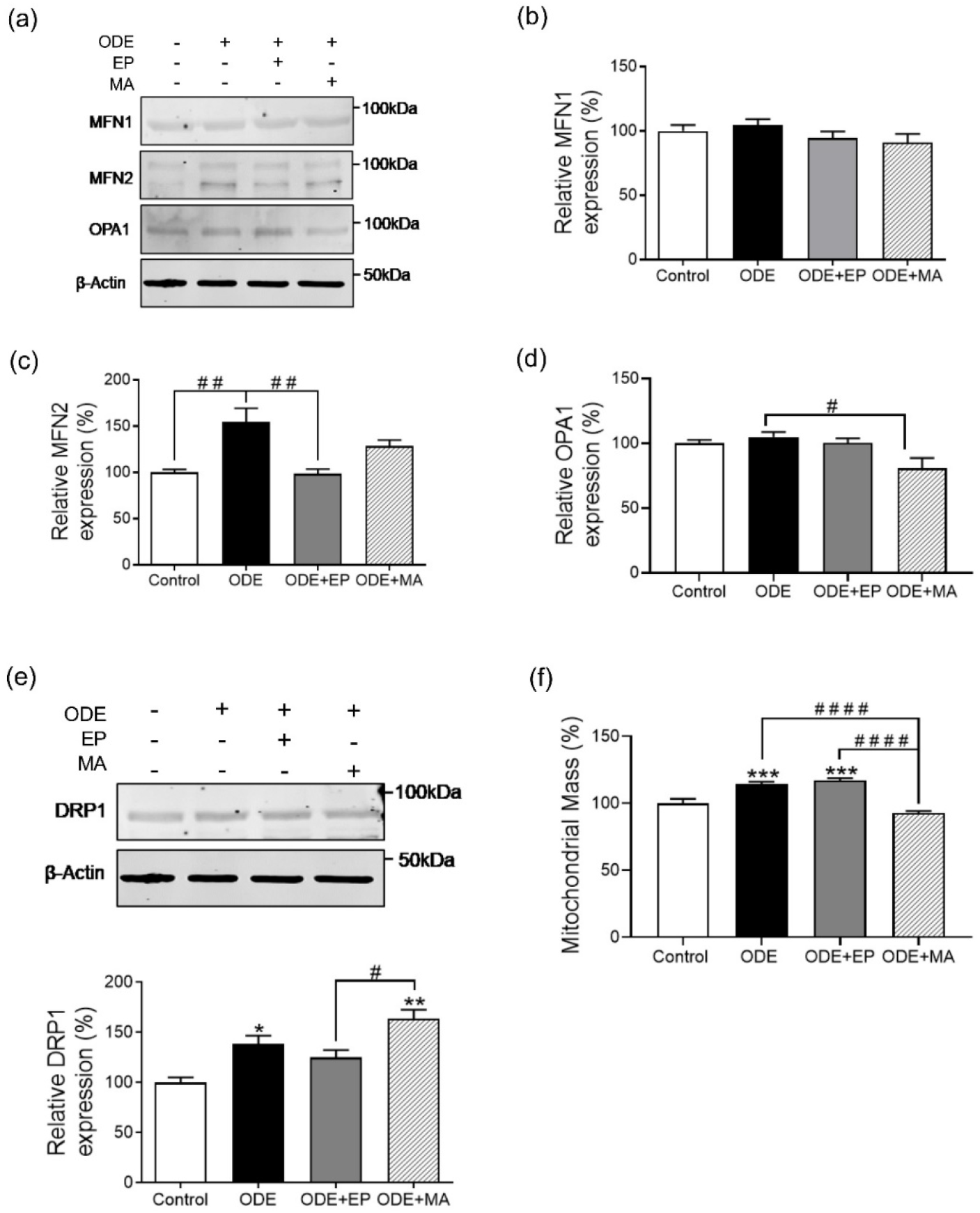
ODE exposure induces fusion of mitochondria in response to stress. Immunoblotting of whole cell lysates of THP1 cells, treated with ODE and antioxidant therapy for 24 hours, was performed to detect mitochondrial fusion and fission proteins. Compared to controls, ODE (1%) treated cells showed minimal changes in the expression of MFN1/2 and OPA1 (a-d), while OPA1 expressions is significantly decreased on exposure to 10 μM of MA. ODE treated cells showed an increase in DRP1 expression, whereas co-treatment with 10 μM of MA, significantly upregulates DRP 1 expression and co-treatment with 2.5 μM of EP downregulates DRP1 comparable to control (e). Mitochondrial mass was measured by Mito-Tracker staining and data showed a significant increase in the mass with ODE (1%) treatment, while 10 μM of MA significantly reduced the mitochondrial mass to the baseline (control) levels (f). For all western blots, samples were derived from the same experiment and were processed in parallel. All protein bands were normalized over β-actin (37 kD) and percentage intensity relative to control analyzed. All data analyzed via one-way ANOVA with Tukey’s multiple comparison test, ^# or^ *p < 0.05, ^# # or^ **p < 0.01, ^# # # or^ ***p < 0.001, ^# # # # or^ ****p < 0.001 and are represented as Mean ± SEM with n = 3-6/treatment.

### ODE exposure induces selective targeting of mitochondria for autophagy (mitophagy)

Due to the increased MFN2 expression observed, it can be questioned whether this increase is favoring the process of mitochondrial elimination (Fig. 3a & 3c). This process involved in the regulation of mitochondrial dynamics is also known to be closely associated with the process of mitochondrial quality control by autophagy, known as mitophagy (Ding and Yin, 2012; Filadi et al., 2018). Mitophagy is critical for maintaining proper cellular functions (Ding and Yin, 2012). Investigation of whether the mitochondria was subjected to autophagic clearance on ODE exposure was done. The expression of the two important mediators of mitophagy, PTEN-induced kinase 1 (PINK1) and the E3 ubiquitin protein ligase Parkin, were investigated. It was observed that there was increased Parkin expression in the presence of ODE compared to controls, while expression of PINK1 remained unchanged (Fig. 4a & 4c). Expression of Parkin remained comparable to the control in the presence of EP or MA (Fig. 4a & 4b). Parkin has been shown to be highly essential in the induction of mitophagy (Ding and Yin, 2012; D. Narendra et al., 2010; Narendra et al., 2008; D. P. Narendra et al., 2010). PINK1 and Parkin are known to physically interact with each other in order to induce mitophagy, and the translocation of Parkin to the mitochondria is said to be dependent on PINK1 (Narendra et al., 2008; D. P. Narendra et al., 2010). This indicates that ODE-induced cellular stress is leading to Parkin mediated mitochondrial clearance.

**Figure 4.**
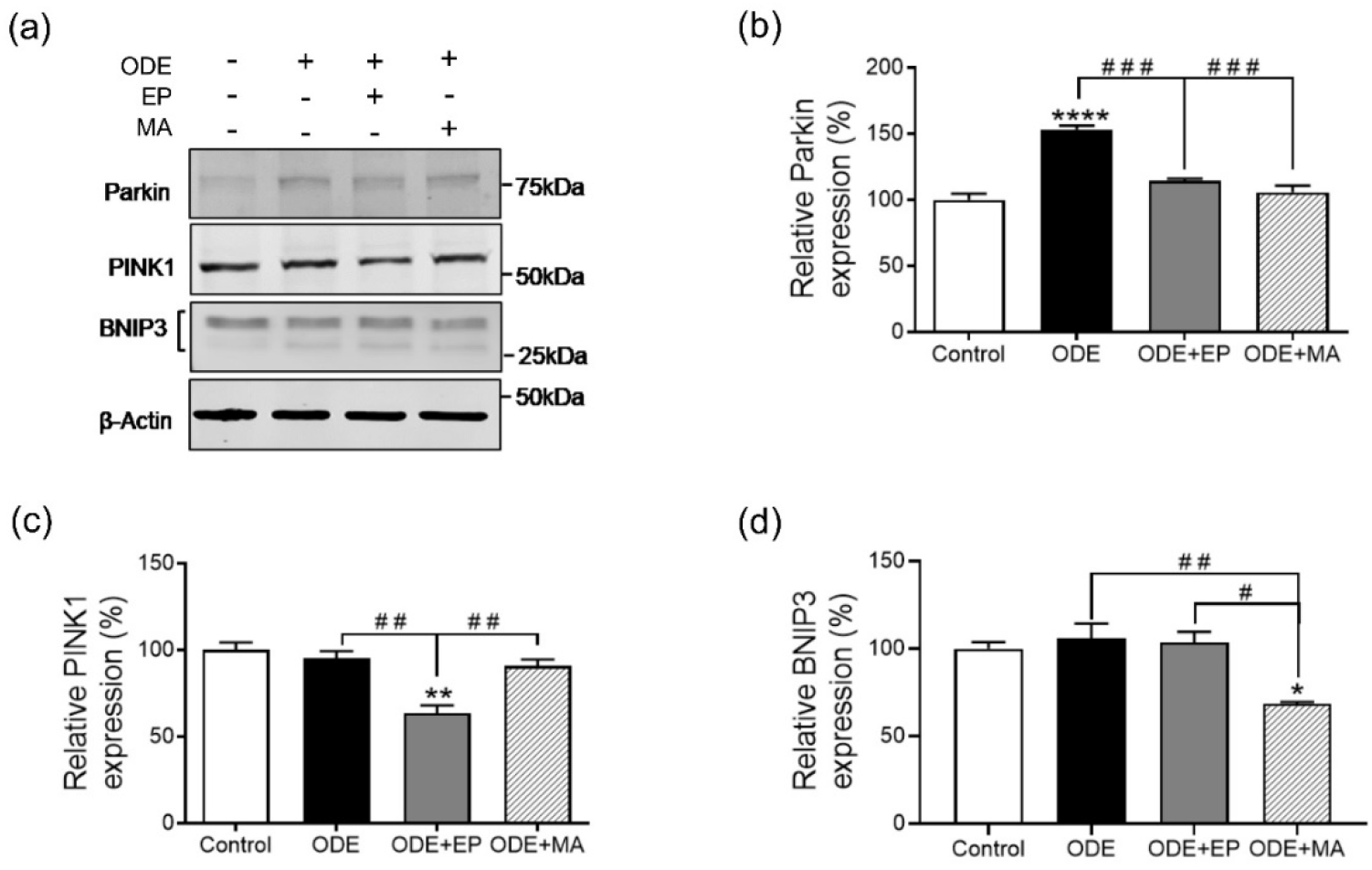
ODE exposure induces selective targeting of mitochondria for autophagy (mitophagy). Immunoblotting of whole cell lysates of THP1 cells, treated with ODE and antioxidant therapy for 24 hours, was performed to detect expression of mitophagy markers. Compared to controls, cells treated with ODE (1%) showed increase in the expression of Parkin (a-b), while expression of PINK1 remains relatively constant (c). Co-treatment with 10 μM of MA significantly decreased Parkin and BNIP3 expressions (b & d). Co-treatment with 2.5 μM of EP significantly decreased PINK1 and Parkin expressions, while having no impact on BNIP3 (a-d). For all western blots, samples were derived from the same experiment and were processed in parallel. All protein bands were normalized over β-actin (37 kD) and percentage intensity relative to control analyzed. All data analyzed via one-way ANOVA with Tukey’s multiple comparison test, ^# or^ *p < 0.05, ^# # or^ **p < 0.01, ^# # # or^ ***p < 0.001, ^# # # # or^ ****p < 0.001 and are represented as Mean ± SEM with n = 3/treatment.

The expression of BNIP3, a mitochondrial Bcl-2 Homology 3 (BH3)-only protein, was also observed. BNIP3 activates the mitochondrial permeability transition (MPT), which is associated with increased ROS production and excessive autophagy (Ney, 2015). BNIP3 levels remained unchanged on exposure to ODE and EP (Fig. 4a & 4d). While, co-treatment with MA significantly decreased BNIP3 expression (Fig. 4d). This indicates that although the process of mitophagy may or may not be occurring via BNIP3, MA is certainly BNIP3 mediated MPT thus having a protective effect on the mitochondria.

### ODE exposure impacts mitochondrial membrane permeability

Mitochondrial oxidative phosphorylation (OXPHOS) pathway is critical in determining and maintaining the immunomodulatory phenotype of activated macrophages (Kelly and O’Neill, 2015). Considering the changes in mitochondrial structure and dynamics, the question of whether ODE has an impact on the mitochondrial OXPHOS pathway was investigated. ODE increased levels of cytochrome c in the cytosol of the cells, compared to that in the mitochondrial fraction (Fig. 5a-5c). Release of cytochrome c is considered a key initial step in the apoptotic process (Cai et al., 1998; Ott et al., 2002). On the other hand, MA significantly decreased the levels of cytosolic cytochrome C (Fig. 5a-c). Concurrently, there is a significant decrease in the expression of lung-specific isoform of cytochrome c oxidase (COX4i2) in the mitochondrial fraction on ODE exposure, with no change on treatment with either EP or MA (Fig. 5d). COX4i2 is considered to be a rate-limiting step of the electron transport chain (ETC) in intact mammalian cells under physiological conditions (Hüttemann et al., 2012). A loss of expression would suggest dysfunctional OXPHOS pathway (Hüttemann et al., 2012). There is also an increase in superoxide dismutase 2 (SOD2) in the cytosol during ODE exposure (Fig. 5e). Treatment with EP or MA significantly decreased SOD2 expression compared to ODE (Fig. 5e). Presence of SOD2 is known to impart tolerance during high oxidative stress and reduce superoxide accumulation withing the mitochondria (Fukui and Zhu, 2010; Ishihara et al., 2015). To identify whether this is true, mitochondrial superoxide levels was measured by MitoSOX fluorescence. ODE significantly decreased the mitochondrial superoxide, while treatment with MA increased the levels comparable to control (Fig. 5f). ODE mediated decrease could either be a consequence of a leaky mitochondrial membrane or the action of high SOD2 expression. Reactive nitrite species (RNS) released into the extracellular environment was measured by griess assay. ODE exposure increased the levels of RNS in media at 24 hours, which treatment with EP and MA significantly attenuated RNS secretion (Fig. 5g). The results collectively show that in response to ODE, there is an increase in mitochondrial membrane permeability leading to leakage of core proteins involved in the maintenance of mitochondrial function. MA, on the other hand, seems to be maintaining the mitochondrial membrane integrity by acting as an inhibitor of peroxynitrite formation and RNS secretion, thus potentially restoring the damage induced on ODE exposure (Ghosh et al., 2016; Stefanska and Pawliczak, 2008).

**Figure 5.**
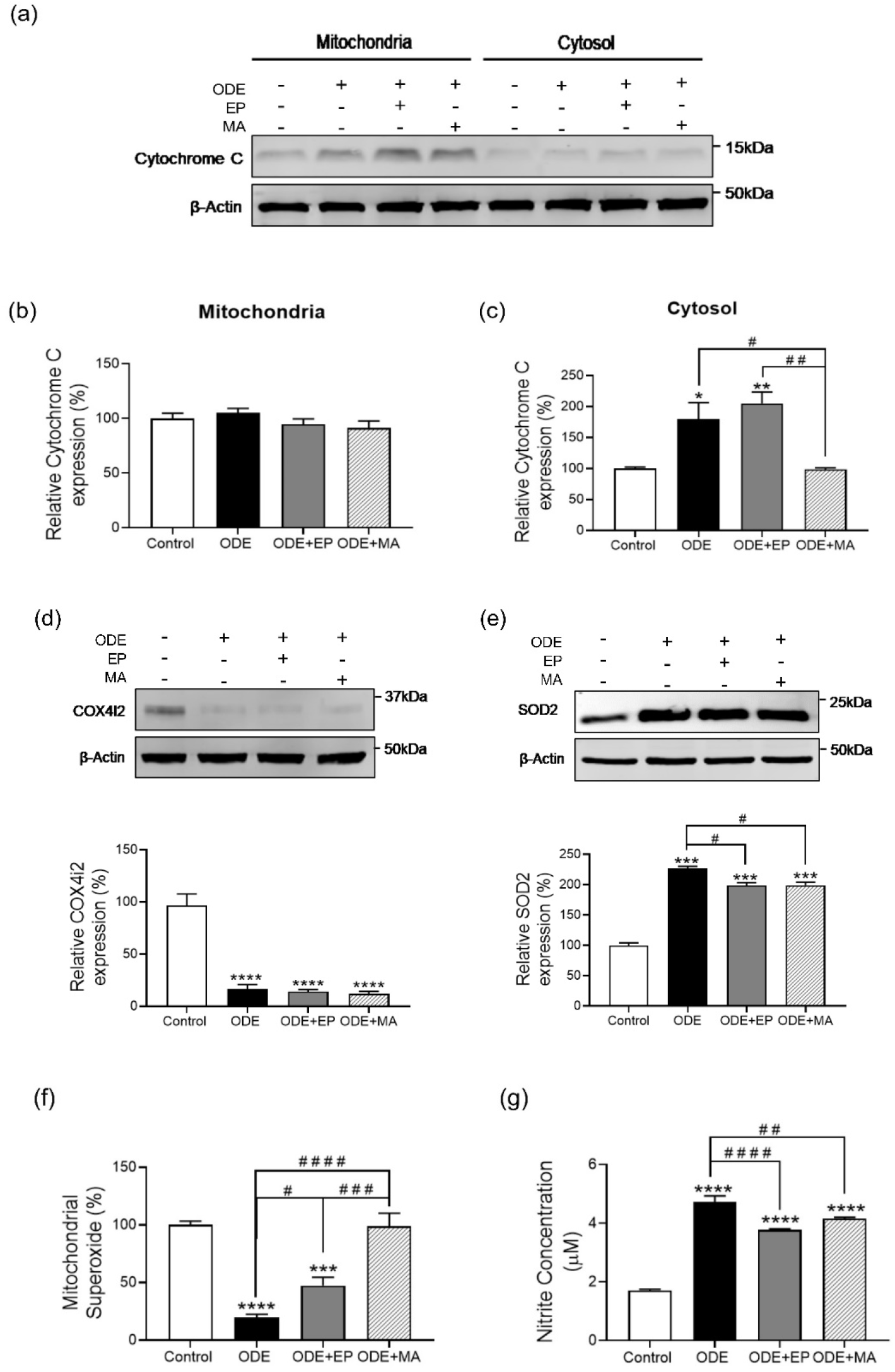
Mitoapocyanin treatment decreases ODE-induced Cytochrome C release and markedly increases SOD2 expression in the cytosol. Immunoblotting of mitochondrial and mitochondria-free cytosolic fractions of THP1 cells, treated with ODE and antioxidant therapy for 24 hours, was performed to detect the presence of Cytochrome C and expression of lung-specific isoform of COX, COX4i2, on ODE exposure. ODE (1%) and 2.5 μM of EP co-treated cells showed a significant increase in Cytochrome C in the cytosol (a & c), while treatment with 10 μM of MA downregulated Cytochrome C comparable to control (c). No change in levels of mitochondrial Cytochrome C was observed for all treatments (b). ODE (1%) treated cells showed a significant decrease in COX4i2 expression (d), while superoxide dismutase 2 (SOD2) increased compared to controls and abrogated when treated with 10 μM of MA and 2.5 μM of EP (e). MitoSOX assay performed showed a decrease in superoxide anions (SOX) on ODE (1%) exposure, while levels on treatment with treated with 10 μM of MA was comparable to control (f). Griess assay was performed to measure the amount of nitrite levels secreted. ODE (1%) treated cells were observed to secrete elevated levels of nitrite (g). Co-treatment with 2.5 μM of EP or 10 μM of MA decreased nitrite levels. For all western blots, samples were derived from the same experiment and were processed in parallel. All protein bands were normalized over β-actin (37 kD) and percentage intensity relative to control analyzed. All data analyzed via one-way ANOVA with Tukey’s multiple comparison test, ^# or^ *p < 0.05, ^# # or^ **p < 0.01, ^# # # or^ ***p < 0.001, ^# # # # or^ ****p < 0.001 and are represented as Mean ± SEM with n = 3-6/treatment.

### ODE induces the secretion of mitochondrial DAMPs

Mitochondrial secondary messengers, which are mitochondrially derived molecules, can act as mitochondrial damage–associated molecular patterns (mtDAMPs) when produced excessively or are secreted into other cellular locations (Cloonan and Choi, 2012). These mtDAMPs result in the induction of a cascade of inflammatory responses withing the cell, thus resulting in adverse effects on the cell and tissue (Zhang et al., 2010). The levels of mitochondrial transcription factor A (mtTFA) measured in the mitochondrial and cytosolic fractions, showed a significant increase in mtTFA in both mitochondria and cytosol on exposure to ODE (Fig. 6a & 6c). While MA treatment increased mtTFA levels in the mitochondria compared to ODE (Fig. 6a & 6b). At normal physiological levels, mtTFA is an important regulator of mitochondrial DNA integrity, which when leaked out from mitochondria acts as a mtDAMP promoting inflammatory responses (Julian et al., 2013). Due to the increase in mitochondrial membrane permeability and dysfunction seen previously, levels of mtDNA leaking into the cytosol was determined. ODE increased cytosolic mtDNA levels, which is abrogated on treatment with EP or MA (Fig. 6d). In addition, there is an increase in calcium (Ca^2+^) influx into the mitochondria on exposure to ODE, with no significant change in the presence of either MA or EP co-treatment compared to ODE (Fig. 6e). An increase in mitochondrial matrix Ca^2+^ levels has been shown to increase ATP production and is a trigger for cell death (Finkel et al., 2015). The expression of mitochondrial HMGB1 was determined, as presence of HMGB1 in the mitochondrial matrix is said to be critical in the regulation of mitochondrial function (Tang et al., 2011). Compared to control, there is a decrease in mitochondrial HMGB1 on exposure to ODE, which does not seem to be rescued in the presence of MA (Fig. 6f). Whereas on EP treatment, mitochondrial HMGB1 is significantly increased compared to both ODE and MA exposure (Fig. 6f). These findings indicate that with the significant impact ODE has on the mitochondrial quality control and biogenesis there is a release of mtDAMPs into the cytosol which could be leading to a cascade of inflammatory responses consequently causing cell death (Qi et al., 2015; Tang et al., 2011).

**Figure 6.**
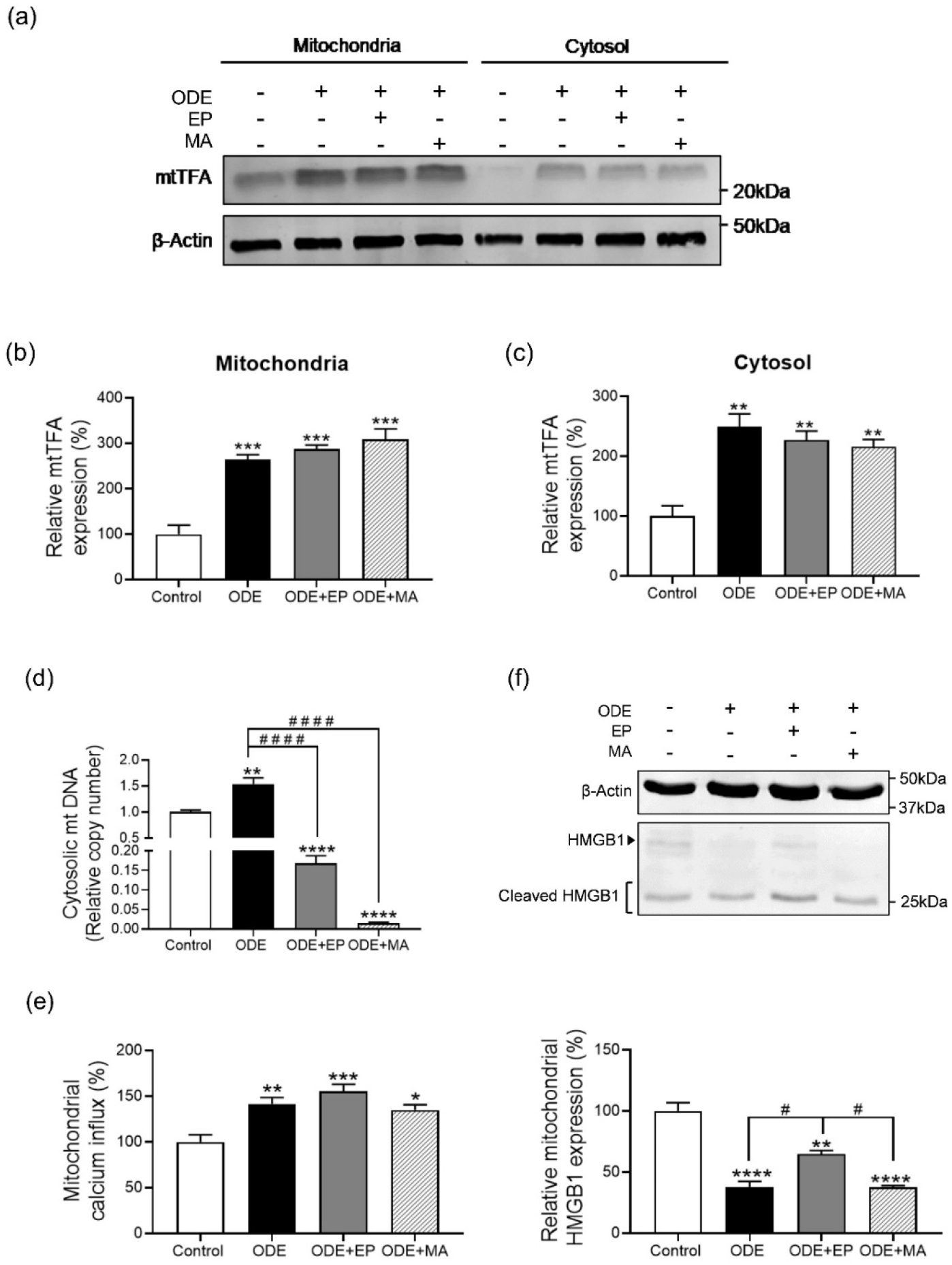
ODE exposure markedly increases secretion of mitochondrial DAMPs into the cytosol. Immunoblotting of mitochondrial and mitochondria-free cytosolic fractions of THP1 cells, treated with ODE and antioxidant therapy for 24 hours, was performed to detect expression of mitochondrial transcription factor A (mtTFA). ODE (1%) treated cells increase mtTFA expression in the cytosol, while treatment with 10 μM of MA increased mtTFA in the mitochondrial matrix (a). Mitochondrial DNA leakage into the cytosol analyzed via qPCR was higher in the cytosol on ODE (1%) exposure which was decreased by 10 μM of MA (d). Intra-mitochondrial calcium levels measured by Rhod 2AM staining, showed increased calcium levels on ODE (1%) exposure, which remained unaffected by 10 μM of MA (e). Immunoblotting of mitochondrial fraction of THP1 cells was performed to measure mitochondrial HMGB1. ODE (1%) treatment showed a decrease in HMGB1, while EP treatment increased the HMGB1 compared to either ODE or MA (f). For all western blots, samples were derived from the same experiment and were processed in parallel. All protein bands were normalized over β-actin (37 kD) and percentage intensity relative to control analyzed. All data analyzed via one-way ANOVA with Tukey’s multiple comparison test, ^# or^ *p < 0.05, ^# # or^ **p < 0.01, ^# # # or^ ***p < 0.001, ^# # # # or^ ****p < 0.001 and are represented as Mean ± SEM with n = 3-6/treatment.

### Mitoapocynin does not intervene in ODE mediated caspase-1 upregulation

As mentioned previously, release of mtDAMPs results in the activation of various inflammatory responses. It has been shown that release of mtDNA and mitochondrial reactive oxygen species (mROS) activates the NLRP3 inflammasome pathway (Gong et al., 2018). Upstream of NLRP3 activation, cleavage of pro-caspase 1 to caspase 1 is seen due to increased influx of calcium induced by leaky mitochondria (Murakami et al., 2012). Based on these evidences, changes in the expression of pro-caspase 1 and caspase 1 was measured. Compared to control, ODE and MA exposure increased expression of pro-caspase 1, EP maintained the expression comparable to control (Fig. 7a & 7b). There was a significant increase expression of cleaved caspase 1 (p10) on ODE exposure and MA co-treatment, with the former inducing a higher expression than the latter (Fig. 7a & 7c). Treatment with EP significantly decreased cleavage, which is consistent with the expression of pro-caspase 1. Expression of pro-caspase 3 and its cleaved product, an apoptosis executioner, was measured in order to determine if ODE is inducing a caspase 3 mediated apoptosis. Although ODE decreased the expression of pro-caspase 3, no significance in the levels was observed on exposure to treatments (Fig. 7d & 7e). In addition, there was no caspase 3 cleavage product observed on exposure to any of the treatments (Fig. 7d). This suggests that ODE could be mediating a downstream inflammatory cascade via caspase 1 cleavage and activation, i.e. NLRP3 inflammasome activation and pro-IL-1β cleavage and release. This is not remedied by co-treatment with either MA or EP. Whereas, caspase 3 may not be a key mediator in ODE mediated inflammation.

**Figure 7.**
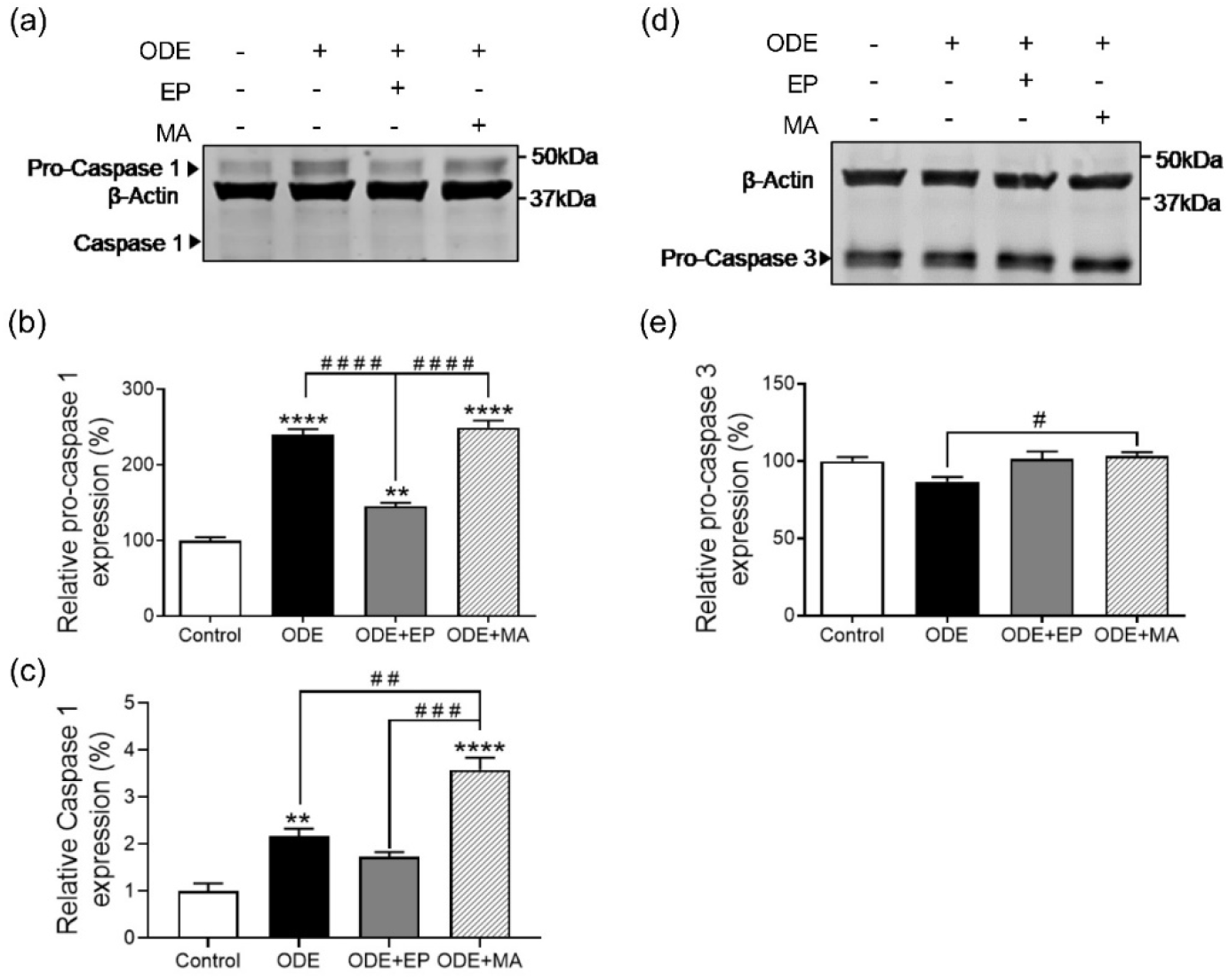
ODE exposure increases expression of Caspase 1 consistent in inflammatory conditions. Immunoblotting of whole cell lysates of THP1 cells, treated with ODE and antioxidant therapy for 24 hours, was performed to detect the expression of caspase 1 and 3. Cells treated with ODE (1%) and 10 μM of MA increase in pro-caspase 1, along with the cleaved caspase 1 p10, while co-treatment with 2.5 μM of EP decreased expression comparable to control (a-c). No significant difference observed with pro-caspase 3 and absence of cleaved caspase 3 (d & e). For all western blots, samples were derived from the same experiment and were processed in parallel. All protein bands were normalized over β-actin (37 kD) and percentage intensity relative to control analyzed. All data analyzed via one-way ANOVA with Tukey’s multiple comparison test, ^# or^ *p < 0.05, ^# # or^ **p < 0.01, ^# # # or^ ***p < 0.001, ^# # # # or^ ****p < 0.001 and are represented as Mean ± SEM with n = 3/treatment.

### Mitoapocynin therapy does not inhibit ODE induced apoptosis

The production of ROS is known to be a trigger for cell death(Brand et al., 2004; Kim, 2005). The antiapoptotic Bcl-2 family proteins Bcl-2 and Bcl-XL play an important role in inhibiting mitochondria-dependent extrinsic and intrinsic cell death pathways (Green and Kroemer, 2004). To identify the impact OD-induced mitochondrial dysfunction and rescue may have on cellular apoptosis, expression of Bcl-2 and Bcl-XL were measured. ODE decreased the expression of Bcl-2, with no change on MA or EP intervention (Fig. 8a). On the other hand, Bcl-XL expression was downregulated on ODE exposure, but was significantly increased on treatment with MA or EP (Fig. 8b). This change in expression of Bcl-XL was corroborated by measuring cell viability by MTT colorimetric assay. The percentage cell viability pattern observed correlated with the patter of expression of Bcl-XL, where loss of cell viability on OD exposure was rescued by treatment with EP or MA (Fig. 8c). Together, these results are indicative that the increase in Bcl-XL expression on treatment with EP or MA, could be blocking the effect of BNIP3 (Fig. 4a & 4d) in inducing the loss of mitochondrial membrane permeability or the activation of caspase dependent or independent apoptotic pathway (Kim, 2005). Thus, regulating the production of ROS and decreasing the probability of apoptotic and non-apoptotic cell death.

**Figure 8.**
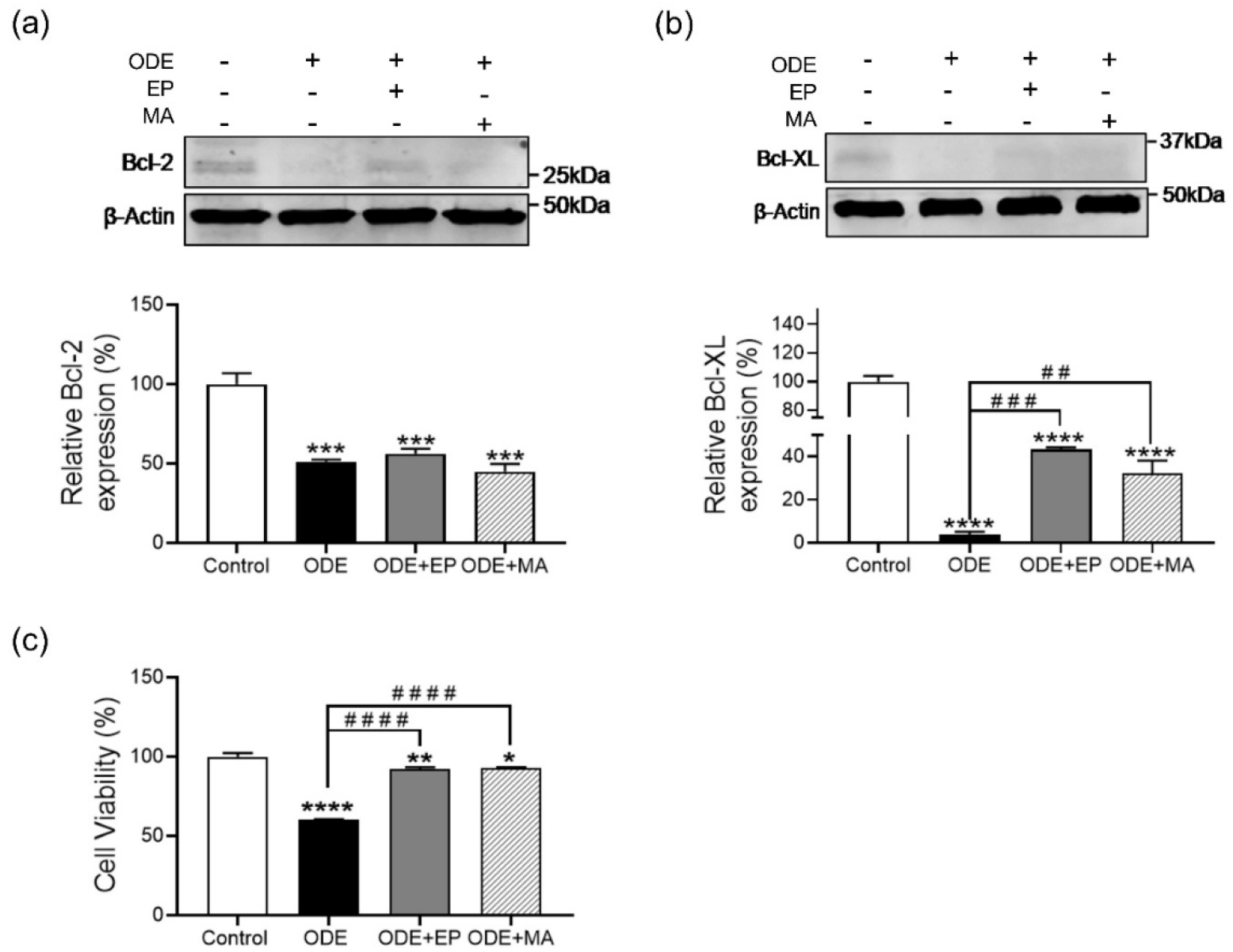
Mitochondrial targeted antioxidant treatment has no effect Bcl-2 and Bcl-XL expression. Immunoblotting of whole cell lysates of THP1 cells, treated with ODE and antioxidant therapy for 24 hours, was performed to observe expression of Bcl-2 and Bcl-XL. ODE (1%) treatment decreased Bcl-2 and Bcl-XL (a-c), while co-treatment with 10 μM of MA increased Bcl-XL comparably higher than ODE (1%; b). MTT assay to measure cell viability showed increased viability on co-treatment with 2.5 μM of EP or 10 μM of MA compared to ODE (c). For all western blots, samples were derived from the same experiment and were processed in parallel. All protein bands were normalized over β-actin (37 kD) and percentage intensity relative to control analyzed. All data analyzed via one-way ANOVA with Tukey’s multiple comparison test, ^# or^ *p < 0.05, ^# # or^ **p < 0.01, ^# # # or^ ***p < 0.001, ^# # # # or^ ****p < 0.001 and are represented as Mean ± SEM with n = 3-6/treatment.

## Discussion

Airway inflammation due to persistent exposure to OD is a key contributor to the development of respiratory symptoms and airflow obstruction in exposed workers (Cole et al., 2000; Nordgren and Charavaryamath, 2018). Continuous exposure to organic dust has been shown to alter innate immune responses in the airways (Charavaryamath and Singh, 2006; Sethi et al., 2017; Wunschel and Poole, 2016). These responses include cellular recruitment, release of pro-inflammatory cytokines and reactive species (ROS/RNS) (Bhat et al., 2019; Nath Neerukonda et al., 2018; Sahlander et al., 2012; Sethi et al., 2017). Previous studies have provided a direct link between such innate immune signaling and mitochondrial dynamics suggesting a crucial role in the activation and control of airway disease progression (Cloonan and Choi, 2012; Eisner et al., 2018). *In vitro* and *in vivo* studies have shown a link between airway diseases such as, influenza, sepsis-induced lung injury, pneumonia, and RSV infection, and facets of mitochondrial responses (Cloonan and Choi, 2016; Wunschel and Poole, 2016). In this study, using THP1 cells as an *in vitro* model for alveolar macrophages we provide evidence of significant changes in the dynamics, integrity and function of cellular mitochondria upon exposure to OD and how the use of mitoapocynin (MA), a novel mitochondrial targeting NOX2 inhibitor, or ethyl pyruvate (EP), an inhibitor of translocation of HMGB1, could rescue OD-induced mitochondrial changes and reduce inflammation.

Our TEM results demonstrate that, upon ODE exposure, there is increased presence of cytoplasmic vacuoles and pseudopods which is a characteristic feature of an activated macrophage ^28^. Treatment with MA or EP did not prevent the ODE-induced morphological changes. When cells were exposed to EP co-treatment, we found formation of multinucleated giant cells (MGC). MGC are a common feature of granulomas that develop during certain infections, the most prominent example being tuberculosis or as a consequence of foreign body reactions (FBR) (Milde et al., 2015; Miron and Bosshardt, 2017).

To understand the impact of OD-induced inflammation on mitochondrial biogenesis, we explored factors involved in mitochondrial morphological changes. Mitochondria are highly dynamic organelles whose functions are essential for cell survival. They continuously change their function, position, and structure to meet the metabolic demands of the cells during homeostatic conditions as well as at times of stress (Eisner et al., 2018; Wai and Langer, 2016). With our TEM findings we observe distinct changes in the mitochondrial surface area and circularity on OD exposure and antioxidant therapy indicating that OD-exposure has an impact on mitochondrial dynamics and functions.

Mitochondria contain outer and inner mitochondrial membranes (OMM and IMM, respectively), which border the intermembrane space (IMS) and the matrix. Each of these compartments has discrete functions in metabolism, biosynthetic pathways, and signaling (Pagliarini and Rutter, 2013). Mitochondrial dynamics involve reshaping, rebuilding, and recycling events that support mitochondrial stability, abundance, distribution, and quality, and allow compensatory changes when cells are challenged (Mishra and Chan, 2014). Key mitochondrial reshaping mechanisms are mitochondrial fission and fusion. Mitochondrial fission is characterized by division of one mitochondrion into two daughter mitochondria, whereas mitochondrial fusion is the union of two mitochondria resulting in one mitochondrion(Mishra and Chan, 2014). The deregulation of these spatio-temporal events results in either a fragmented network characterized by a large number of small round-shaped mitochondria or a hyper fused network with elongated and highly connected mitochondria (Tilokani et al., 2018; Wai and Langer, 2016). These balanced dynamic transitions are not only required to ensure mitochondrial function but also to respond to cellular needs by adapting to nutrient availability and metabolic state of the cell.

Mitochondrial fusion is mediated by dynamin-related GTPases mitofusin 1 and 2 (MFN1/2) on the outer mitochondrial membrane (OMM) and by dynamin-related protein optic atrophy 1 (OPA1) on the inner mitochondrial membrane (IMM). Mitochondrial fission requires the recruitment of dynamin-related protein 1 (DRP1) from the cytosol to its specific receptors (Mishra and Chan, 2014; Tilokani et al., 2018). Lack of either MFN1 or MFN2 expression can display aberrant mitochondrial morphology. While a lack of MFN1 induces mitochondrial fragmentation, the absence of MFN2 exhibits swollen spherical mitochondria. In our findings, we see increased MFN2 expression on OD exposure, which could be leading to increased mitochondrial fragmentation and increased mitochondrial mass (Chen et al., 2003). An increase in the rate of mitochondrial proliferation is probably a cellular response to counteract the loss of mitochondrial function and recover ATP synthesis capacity. We also see an increase in DRP1 expression upon exposure to both ODE and MA co-treatment. Mitochondrial fission is essential for the inheritance and partitioning of mitochondria during cell division. Inhibition of DRP1-mediated mitochondrial fission has been reported to cause cellular dysfunction and replication (Qi et al., 2015). This can be corroborated by the decrease in cell viability with ODE exposure. The low mitochondrial mass observed with exposure to MA could be a means by which the antioxidant therapy is overcoming the increase in dysfunctional mitochondria.

Studies have shown that the MFN2 mediates Parkin, an E3 ubiquitin ligase, recruitment to damaged mitochondria (Filadi et al., 2018). Parkin binds to MFN2 in a PINK1-dependent manner and promotes Parkin-mediated ubiquitination of damaged mitochondria, thus leading to a mitochondria quality control process, known as mitophagy (Ding and Yin, 2012; Narendra et al., 2008; D. P. Narendra et al., 2010). This corroborates our finding wherein we see an increase in MFN2 and Parkin expression upon ODE exposure, from which we can conclude that OD-induced cell stress leads to mitophagy. Mitophagy is a process of mitochondrial quality control where damaged or defective mitochondria are removed by selective encapsulation into double-membraned autophagosomes that are delivered to lysosome for degradation (Ding and Yin, 2012). Mitochondrial biogenesis and mitophagy allow cells to quickly replace metabolically dysfunctional mitochondria.

Albeit no significant change in BNIP3 expression was observed with exposure to OD or EP. However, BNIP3 expression was decreased on intervention with MA. This brings into question how the low levels of BNIP3 is affecting the mitochondrial and cellular function. BNIP3, a transmembrane protein located in the OMM, imparts some pro-cell death activity and is known to regulate mitophagy (Ney, 2015). BNIP3 has been shown to activate the mitochondrial permeability transition (MPT) and degradation of proteins involved in oxidative phosphorylation, in turn leading to cell death without cytochrome c release or caspase activation (Landes et al., 2010; Quinsay et al., 2010; Rikka et al., 2011; Velde et al., 2000). On addition of BNIP3 to isolated mitochondria, it was observed that BNIP3 caused cytochrome c release, depolarization, and swelling (Kim et al., 2002). This phenomenon has been linked to BNIP3-mediated permeabilization of inner and outer mitochondrial membrane involving the disruption of OPA1 complex and remodeling of the inner mitochondrial membrane (Landes et al., 2010). In comparison to our findings, we can assume that MA-induced decrease in BNIP3 expression could be reducing MPT and cell death, thus improving overall cellular function. Another potential mechanism by which BNIP3 is promoting apoptosis is by competition for binding to Bcl-2 (or a related protein) which liberates Beclin-1 from Bcl-2 complexes and activates autophagy. There is evidence showing that Bcl-XL enhances BNIP3-induced mitophagy (Maiuri et al., 2007; Pattingre et al., 2005). This correlates with our findings wherein we see decreased Bcl-XL expression on exposure to OD which is significantly upregulated on intervention with EP or MA. Taken together we see a decrease in overall cell viability on OD exposure which is rescued by MA intervention.

One of the prominent players in cell death is cytochrome c. Based on previous studies, we expected that with the decrease in cell viability and induction of mitophagy there would be a release of cytochrome c from the mitochondria into the cytosol. Cytochrome c, a peripheral protein of the mitochondrial inner membrane (IMM), is known to function as an electron shuttle between complex III and complex IV of the respiratory chain and its activity (Cai et al., 1998; Garrido et al., 2006). And its release from the IMM has been implicated in caspase activation and mitochondrial outer membrane permeabilization (MOMP), leading to cell death. Cumulative data suggest that cytochrome c release does not always take place in an all-or-nothing manner as previously believed, but instead follows a biphasic kinetics (Ott et al., 2002). The first wave is induced by apoptotic signals directed to mitochondria which provokes MOMP and cytochrome c release, thus disrupting the electron transport and leading to an increased generation of ROS. The second wave involves cytochrome c mediated activation of caspases that subsequently enters the mitochondria through the permeabilized OMM and induce the complete block of the respiratory chain, eventually resulting in cell death. Comparing this to our findings we observe that with the increase in cytosolic cytochrome c on OD exposure there is a deficiency of levels of COX4i2 (COX subunit 4 isoform 2), a terminal enzyme in the OXPHOS machinery. Loss of COX4i2 results in decreased COX activity and decreased ATP levels (Hüttemann et al., 2012). This loss is not reversed by the use of antioxidant therapy, albeit MA was capable of downregulating the release of cytochrome c. This is indicative that although antioxidant therapy can decrease the cytosolic release of cytochrome c, there could be other secondary factors resulting in the loss of COX4i2. NADPH oxidase is the main source of ROS that is closely linked to mitochondrial ROS production (Zorov et al., 2014). Growing evidence suggests that ROS generated can increase expression of proinflammatory mediators (Bhat et al., 2019; Brand et al., 2004). Indeed, various PAMP molecules can stimulate ROS production by NADPH oxidase, especially NOX1, NOX2, NOX4 (Ghosh et al., 2016). Being a NOX2 inhibitor, treatment with MA seems to bring mitochondrial superoxide levels to that of the controls, whereas with OD and EP we see a decrease. This could be a consequence of a leaky mitochondria which is enabling the release of the superoxide ion into the cytosol thus promoting further damage to the cell. On the other hand, seeing the increase in the SOD2 expression allows us to believe that there are factors promoting the attenuation of oxidative stress mediated cellular injury. This increase could be due to a variety of proinflammatory cytokines, such as interleukin 1 (IL-1), IL-4, IL-6, tumor necrosis factor α, interferon γ, and the bacterial endotoxin lipopolysaccharide, which are considered to be robust SOD2 activators (Fukui and Zhu, 2010, p. 2). SOD2 is also said to be regulated by RNS, where increased peroxynitrite levels can lead to its enzymatic inhibition (Redondo-Horcajo et al., 2010). These antagonistic roles that peroxynitrite and superoxide radicals have in regulating SOD2 expression and activity leads us to believe that mitochondrial antioxidant response is dysregulated.

A wide variety of mitochondrial-derived molecules, which act as second messengers, can also behave as mitochondrial damage–associated molecular patterns (mtDAMPs) when produced in excess or released into an alternative cellular compartment. Activation of MPT during mitochondrial dysfunction has also been shown to cause leakage of mtDAMPs, primarily mitochondrial DNA (mtDNA) into the cytosol and activating caspase 1 (Nakahira et al., 2011). The release of mtDNA has been shown to cause neutrophil mediated organ injury by systemic inflammatory reaction via the activation of DNA sensor cyclic GMP–AMP synthase (cGAS) and TLR9 pathway, intracellularly (West et al., 2015; Zhang et al., 2010). Mitochondrial transcription factor A (mtTFA) is an integral regulator of mtDNA integrity, which, when released from mitochondria, acts as a mtDAMP to regulate inflammatory responses (Julian et al., 2013). Release of mtTFA along with mtDNA during cell damage amplified TNFa and type 1 interferon release, which plays a critical role in promoting sterile inflammation and autoimmune diseases (Cantaert et al., 2010; CHAUNG et al., 2012; Julian et al., 2012). This is in line with our findings where we see an increase in the cytosolic release of mtTFA and mtDNA on OD exposure. Although antioxidant therapy did not have a significant impact on reducing release of mtTFA, it did however decrease the release of mtDNA. Being a homolog of mtTFA, HMGB1 was investigated to understand the impact of its translocation into the mitochondrial can have (Parisi and Clayton, 1991). Under pathophysiological conditions, nuclear HMGB1 is immediately transported to the cytoplasm and released into the extracellular space where it acts as a signaling molecule regulating a wide range of inflammatory responses by binding to TLR2/4 and/or receptor for advanced glycan end products (RAGE) (Bhat et al., 2019). It has been reported that HMGB1 rescues the impairment of mitochondrial function. In endothelial cells, the translocation of endogenous HMGB1 from the nucleus to the mitochondria promotes mitochondrial reorganization (Hyun et al., 2016; Stumbo et al., 2008). In cancer cells, exogenous HMGB1 enters the mitochondria, which is followed by the formation of giant mitochondria (Gdynia et al., 2016; Hyun et al., 2016). Therefore, it is likely that the nuclear HMGB1 export would be involved in aberrant mitochondrial fission or the compensatory responses for maintenance of mitochondrial functions. However, in the present study we see that treatment with MA does not revert back the levels of HMGB1 within the mitochondria and match the levels observed on OD exposure.

Mitochondria are also key regulators calcium (Ca^2+^) which control a diverse range of cellular processes, including ROS production. Ca^2+^ influx into the mitochondrial matrix ([Ca^2+^]_mito_) has been shown to be an important regulator of mitochondrial metabolism, and the mobilization of and regulation of mitochondrial Ca^2+^ uptake has been linked to Bcl-XL and voltage-dependent anion channel (VDAC) interactions (Huang et al., 2013; Jouaville et al., 1999; Pitter et al., 2002). Any aberrant increase in cytosolic Ca^2+^ and resultant [Ca^2+^]_mito_ overload can be a trigger for cell death (Finkel et al., 2015). This overload has also been linked to induction of MPT, resulting in mitochondrial permeabilization (Finkel et al., 2015; Hunter et al., 1976). This is corroborated by our findings where we see increase in expression of MPT-inducing markers and [Ca^2+^]_mito_ levels on OD exposure. However, on our observation that Bcl-XL is abrogated, it is safe to assume that the Ca^2+^ influx could possibly be occurring via the interaction of VDAC with Mcl-1, a Bcl-2 family protein (Huang et al., 2014). Ca^2+^ signaling also plays a critical role in the activation of NLRP3 inflammasome by multiple stimuli (Murakami et al., 2012). This is corroborated by our findings wherein we see caspase-1 processing with the increase Ca^2+^ levels on OD exposure (Yu et al., 2014). This would in turn lead to IL-1β processing and release into the extracellular space. The use of MA does not seem to have any impact on the levels of Ca^2+^ accumulation within the mitochondria which could be inducing an inflammatory cascade not mediated by mitochondria.

## Conclusion

In conclusion, we document that co-treatment with EP and MA are partially protective as they rescue some of the ODE-exposure induced deficits. However, these findings lead to new mechanistic questions on how OD may be inducing mitochondrial dysfunction and cell death. OD being a complex mixture of contaminants could be inducing a multifactorial immune response and the mechanism underlying these responses are not yet well understood. Specific signatures of mitochondrial dysfunction that are associated with disease pathogenesis and/or progression are becoming increasingly important. Although our current study is limited with the use of a single immortalized cell line as a model, it provides data on the impact of OD on mitochondrial biogenesis and function. Future studies using functional (primary alveolar macrophages, precision cut lung slices) and mouse model would be highly valuable.

## Acknowledgements

We would like to thank Tracey Stewart at ISU’s Roy J. Carver High Resolution Microscopy Facility for assistance with transmission electron microscopy.

## Author contributions

S.M. Bhat participated in the design of experiments, performed the experiments, analyzed the data, and wrote the manuscript. D. Shrestha performed the calcium influx assay. N. Massey performed organic dust extraction. L. Karriker collected the organic dust samples and edited the manuscript. A.G. Kanthasamy provided mitoapocynin and edited the manuscript. C. Charavaryamath conceptualized the study, participated in the design of the experiments, performed dust extraction, participated in the interpretation of data and edited the manuscript. All authors have read and approved the final manuscript.

## Abbreviations

OD: Organic Dust
ODE: Orgaic Dust Extract
EP: Ethyl Pyruvate
MA: Mitoapocynin
LPS: Lipopolysaccharide
PGN: Peptidoglycan
PAMPs: Pathogen Associated Molecular Patterns
COPD: Chronic Obstructive Pulmonary Disease
AHR: Airway hyperresponsiveness
ROS: Reactive Oxygen Species
RNS: Reactive Nitrogen Species
ATP: Adenosine Triphosphate
OXPHOS: Oxidative Phosphorylation
HMGB1: High Mobility Group Box 1
STAT: Signal Transducer and Activator of Transcription
TPP: Triphenylphosphonium
MPTP: 1-Methyl-4-Phenyl-1,2,3,6-Tetrahydropyridine
iNOS: inducible Nitric Oxide Synthase
NOX: NADPH Oxidase
MTT: 3-[4,5-dimethylthiazole-2-yl]-2,5-diphenyltetrazolium bromide
TEM: Transmission Electron Microscopy
DMSO: Dimethyl Sufoxide
mtND1: mitochondrial NADH dehydrogenase 1
MFN: Mitofusin
OPA1: Optic Atrophy 1
DRP1: Dynamin-related protein 1
ER: Endoplasmic Reticulum
PINK1: PTEN- induced kinase 1
BNIP3: Bcl-2 Homology 3 (BH3)-only
MPT: Mitochondrial Permeability Transition
COX4i2: Cytochrome C Oxidase subunit 4 isoform 2
ETC: Electron Transport Chain
SOD2: Superoxide Dismutase 2
mtDAMPs: mitochondrial Damage Associated Molecular Patters
mtTFA: mitochondrial Transcription Factor A
MGC: Multinucleated Giant Cell
FBR: Foreign Body Reactions
OMM: Outer Mitochondrial Membrane
IMM: Inner Mitochondrial Membrane
IMS: Intermembrane Space
IL: Interleukin
cGAS: cyclic GMP-AMP synthase
TLR: Toll-like receptor
RAGE: Receptor for advanced glycation end products
VDAC: Voltage-dependent anion channel

